# Dynamic vs. Static Facial Color Changes: Evidence for Terminal Color Dominance in Expression Recognition

**DOI:** 10.1101/2025.03.06.641772

**Authors:** Miku Shibusawa, Yuya Hasegawa, Hideki Tamura, Shigeki Nakauchi, Tetsuto Minami

## Abstract

Facial color is closely linked to the perception of emotion, with reddish tones often associated with anger. While previous studies have shown that static reddish facial tones enhance this perception, it remains unclear whether dynamic changes in facial color further amplify this effect. This study investigated how variations in facial color influence the perception of expression using a judgment task that involved morphed facial stimuli (fearful to angry). The participants evaluated facial expressions under two conditions: faces with dynamic color changes and faces with static colors. Experiment 1 compared red (CIELAB a*+) faces to original-colored faces, and Experiment 2 compared green (a*−) faces to original-colored faces. Neither experiment revealed significant differences between dynamic and static facial colors. However, faces with a final reddish color (higher a* value) were more likely to be perceived as angry. These findings suggest that the final facial color influences the perception of anger independent of whether the color change is dynamic or static. Our findings support the idea that the recognition of anger is modulated by the relationship between an angry expression and the color red and provide a new perspective on facial color changes in the interaction between facial expression and facial color.

## Introduction

Human faces undergo dynamic changes, and dynamic changes to facial expressions facilitate the perception of emotion (Ambadar et al., 2005; Fujimura & Suzuki, 2010; Krumhuber et al., 2013; Recio et al., 2011; Trautmann et al., 2009). For example, it has been reported that stimuli that involve dynamic changes in facial expressions are related to greater accuracy in the judgment and categorization of emotion than static facial expression stimuli (Ambadar et al., 2005; Fujimura & Suzuki, 2010). In addition to facial expressions, facial color changes depending on emotion (Drummond et al., 2001; Kreibig, 2010). For example, physiological responses such as heart rate, blood pressure, and skin temperature change in relation to emotional states (Kreibig, 2010). Additionally, facial blood flow increases when people express anger, leading to facial redness (Drummond et al., 2001).

Facial color is closely related to emotion, as in the relationship between the color red and anger, and this relationship may affect a person’s recognition of facial expressions (Hasegawa et al., 2025; Jonauskaite et al., 2020; Liao et al., 2018; Nakajima et al., 2017; Nguyen et al., 2023; Qin, 2021; Takahashi & Kawabata, 2018; Takei & Imaizumi, 2022; Thorstenson et al., 2018; Thorstenson, McPhetres, et al., 2021; Thorstenson & Pazda, 2021). In the judgment of facial expressions, a reddish face is likely to be perceived as angry even if the shape of the face remains the same (Nakajima et al., 2017). In particular, red has been reported to influence the perception of angry facial expressions even under strict conditions that control for facial color changes and the number of emotions and colors (Peromaa & Olkkonen, 2019). From a physiological perspective, Kato et al. (2022) reported that facial stimuli with increased color, which is associated with hemoglobin concentration, increased the perception of angry faces (Kato et al., 2022). Moreover, this effect of increased emotion based on color occurs not only in real faces and facial colors but also in model faces and background colors (Minami et al., 2018; Qin, 2021; Thorstenson, McPhetres, et al., 2021). In contrast to the enhanced perception of anger due to a reddish facial color, it has been reported that green, the color opposite red, reduces the perception of anger in the classification of disgust versus anger (Thorstenson, McPhetres, et al., 2021). Moreover, reddish angry faces increase the perception of emotional intensity, aggression and threat, whereas greenish angry faces decrease these perceptions (Thorstenson, Pazda, et al., 2021; Thorstenson & Pazda, 2021). Although facial color modulates people’s response to facial expressions, cases have been reported in which facial expressions influence facial color. For example, recall and memory of facial color shift toward warmer colors (a*+ and b*+) for angry faces, while the perception of facial color shifts toward blue (b*−) for sad faces (Hasegawa et al., 2024; Nakajima et al., 2017; Thorstenson, Pazda, et al., 2021). Thus, the relationship between facial expressions and color, such as angry faces and the color red, affects human perceptions and responses.

Although facial changes are dynamic in nature, static facial color stimuli have been used in many previous experiments (e.g., Nakajima et al., 2017). Most studies that incorporate dynamic changes examine changes to both facial expression and color together or focus solely on dynamic changes to facial expression. Thorstenson et al. (2021) reported that facial stimuli with dynamic changes to expression and color generally facilitated more accurate categorization of emotion compared with the absence of changes to facial color (Thorstenson, Pazda, et al., 2021). However, that study did not compare dynamic changes to facial color with pre-colored changes to facial color (i.e., facial colors that are consistently reddish or yellowish) but investigated only the effects of facial coloration. Therefore, the influence of dynamic changes to facial color alone on the recognition of facial expression remains unclear. For example, the judgment of facial expression depends on the color of the face, but it is uncertain whether this effect differs if the facial color is dynamic or static. Nakajima et al. (2017) reported that when participants judge the facial expressions of morphed facial image stimuli ranging from fear to anger, reddish faces increase the perception of anger, even at the same morphing level. Is this increased perception of anger further enhanced by dynamic changes in facial coloration?

This study aimed to clarify the effect of dynamic changes to facial color on the identification of facial expressions. We conducted a facial expression judgment task with the following hypothesis: if dynamic changes to facial color bias people’s recognition of facial expressions more than static facial color does, the perception of anger will be stronger when facial color changes dynamically from neutral to red than when facial color is consistently red. On the basis of a study by Nakajima et al. (2017), we conducted experiments in which the participants selected the emotion of a morphed facial expression stimulus with dynamic or static facial color from two choices and compared the level of facial perception between the facial color conditions (Nakajima et al., 2017). When the focus is only the effect of changes in facial color, facial expressions remain static.

## Experiment 1

We conducted a facial expression judgment task to investigate the effects of differences in facial color and the presence or absence of changes in facial color on the recognition of facial expressions following the methods outlined by Nakajima et al. (2017).

### Participants

Twenty-two Japanese students (9 women and 13 men; average age 21.6±1.05 years, between 18 and 24 years old) at Toyohashi University of Technology participated in Experiment 1. The sample size was calculated using PANGEA with an effect size of *d* = 0.5, *α* = 0.05 and *power* > 0.8 (Westfall et al., 2014). We used the Ishihara Color Vision Test Chart II Concise Version 14 Table (Public Interest Incorporated Foundation Isshinkai, Handaya Co., Ltd., Tokyo, Japan) to check the participants’ color vision. All participants had normal color vision as verified by the test chart. They were provided with an introduction to the experiment (excluding the study’s hypothesis) and provided informed consent to participate. This experiment was conducted with the approval of the Ethics Committee for Human Research at Toyohashi University of Technology.

### Stimuli

An eleven-step morphing of facial images from fearful (0) to angry expressions (10), as in a previous study (1 woman, 1 man; Nakajima et al., 2017), was used in this experiment. The facial color of each image was changed across the four conditions (see Figure 1). In the facial color change conditions, the color was changed from original (a*: ±0) to red (+12) or red to original in the CIEL*a*b* color space. The facial color was changed linearly over the course of 1 second, and each condition was defined as “OR (original to red)” or “RO”. To compare the facial color change and no-change conditions, we provided the original (±0) and red (+12) facial colors in the no-change condition. The facial color did not change during the presentation under these conditions. We defined these conditions as “OO (original to original = consistently original)” and “RR”. The changing value and time were determined on the basis of previous research (Nakajima et al., 2017; Thorstenson, Pazda, et al., 2021). The size of the stimuli was 3.14° × 3.14°, and the background color was always gray (*Y* = 21.0 cd/m^2^).

**Figure 1.**
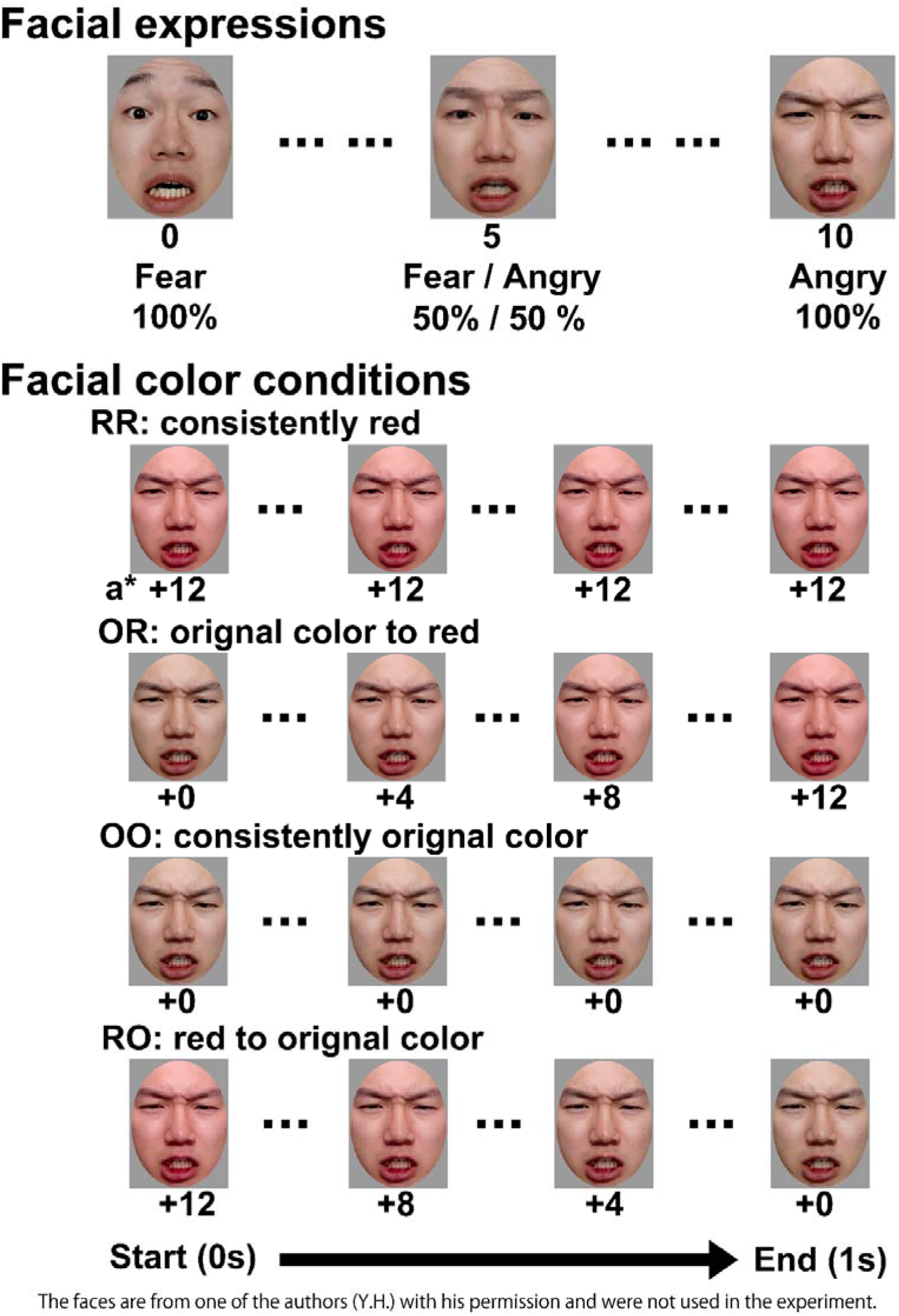
Examples of the image stimuli for each condition in Experiment 1. An 11-step (0: most fearful to 10: most angry) morph of facial stimuli was used. See Nakajima et al. (2017) for more specific morphing image stimuli(Nakajima et al., 2017). Four facial color conditions were provided using red and the original color. The faces in the figure are from one of the authors (Y.H.) with his permission and were not used in the experiment.

### Apparatus

The experiment was conducted in a dark booth. The stimuli were presented on a monitor (EIZO CG319X; EIZO Corporation, Hakusan, Ishikawa, Japan; resolution: 1920 × 1080; frame rate: 60 Hz) that was calibrated with SpyederX Elit (Datacolor, Lawrenceville, NJ, USA) and ColorNavigator 7 (color management software provided by EIZO). The white point chromaticity of the monitor was *x* = 0.30, *y* = 0.33, *Y* = 92.2 cd/m^2^. The participants were seated and performed the task while keeping their heads on a chin rest positioned 85 cm from the display. MATLAB R2024a and Psychotoolbox 3.0.17, toolboxe of MATLAB, served as the experimental control software(Brainard, 1997; Kleiner et al., 2007; Pelli, 1997).

### Procedure and task

On the basis of previous studies (Nakajima et al., 2017), we conducted a facial expression judgment task. Figure 2 shows a summary of the experimental procedure. After a 0.5 second interstimulus interval and 1.0 second fixation were presented, the participants viewed a facial stimulus for 1.0 second. Then, colorful noise was provided for 1.0 s to reduce the aftereffects to the phase of facial expression judgment. The participants identified the facial expression of the presented stimulus as “anger” or “fear” via a numeric keypad. The participants were instructed to respond intuitively and to prioritize accuracy. There were 88 facial stimuli (11-step morphed faces x 2 individuals x 4 facial color conditions), and each stimulus was presented 5 times in random order (total of 440 trials per participant). The experiment consisted of 44 trials in one block (total of 10 blocks). The participants could take breaks between blocks.

**Figure 2.**
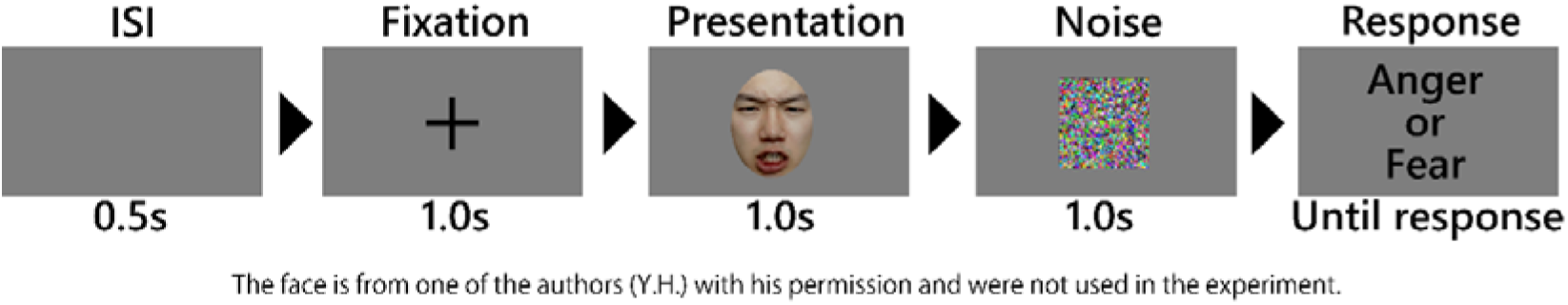
Procedure of the experiment (Experiment 1 & 2). The facial color changed during the presentation phase (1.0 second) in the facial color change conditions. Notably, the ratio of the screen to the fixation cross, facial stimulus, noise image, and text in this figure differs from the actual ratio.

Before the main session, we conducted a practice session to introduce the facial stimuli with angry or fearful expressions. In the practice session, we used the angriest (10) and the most fearful face (0) for each gender. The facial color of the stimuli was consistently the original color (as in the OO condition), and each stimulus was assessed twice (for a total of 8 trials). Feedback on correctness or incorrectness was presented on the monitor after the response only in the practice session. The practice session was established on the basis of the results of preliminary experiments, and the results were not used in the analysis.

### Data analysis

In each condition, the rates of identification of each facial expression were fitted for each participant with a psychometric function. We used a generalized linear model with a binominal distribution in the MATLAB Palamedes Toolbox as the psychometric function (Prins, 2013, 2023; Prins & Kingdom, 2018). Before the fitting, we excluded poorly performing participants whose identification rates in the 0 (the easiest fearful face) and 10 (the easiest angry face) conditions were outside the plus and minus three sigma ranges of the participants’ averages, respectively. As a result, three participants were excluded from the analysis. Nineteen participants’ data were used in Experiment 1. In each condition, the point of subjective equality (PSE) for each participant was calculated from the psychometric curve. The PSE was the morphed level of facial expression at which the probability of identifying the facial expression was equal for angry and fearful faces. We compared the PSEs of each condition to investigate whether they differed by the presented facial color.

In the statistical analysis, we first conducted the ShapirolJWilk test and confirmed that the PSEs of each condition were normally distributed. We subsequently performed a repeated-measure one-way analysis of variance using R (version 4.4.1). Additionally, the *p* values were adjusted using the Holm method in the post hoc tests.

### Results and discussion

Figure 3 shows the mean PSE for each facial color condition. We found significant differences between the facial color conditions (*F*(1.35,24.26) = 7.57, *p* = .007, *η_p_*^2^= .296). The post hoc test results revealed that the PSEs for OR and RR were smaller than those for OO (OR vs. OO: *t*(18) = 3.05, *adj. p* = .035, Cohen’s *d* = 0.70; RR vs. OO: *t*(18) = 3.87, *adj. p* = .007, Cohen’s *d* = 0.89). However, there were no significant differences between OR and RR or between RO and OO (OR vs. RR: *t*(18) = 0.22, *adj. p* > .999, Cohen’s *d* = 0.05; RO vs. OO: *t*(18) = 0.51, *adj. p* > .999, Cohen’s *d* = 0.12). The other results of the statistical analysis for the post hoc test are shown in Table 1.

**Figure 3.**
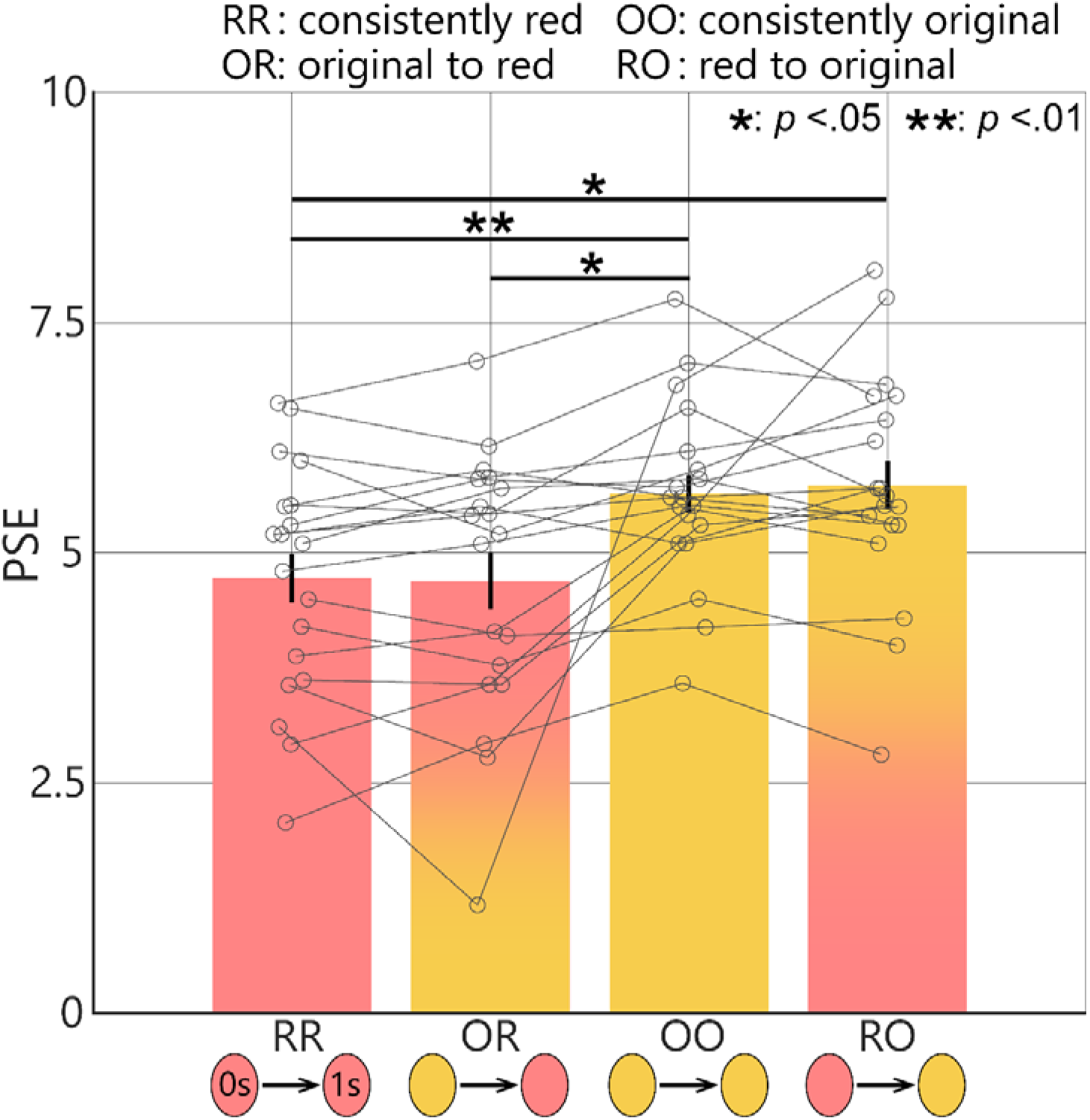
Participants’ mean of PSE in Experiment 1. A smaller PSE value indicates that the participants were more likely to perceive faces in that condition as angry faces, whereas a larger value indicates that the participants were more likely to perceive faces in that condition as fearful faces. The points and lines of each gray circle represent individual data, and error bars indicate the standard error of the mean. The color below each bar in the bar graph represents the facial color at the beginning of the presentation (0 s), while the color above each bar represents the facial color at the end of the presentation (1 s).

**Table 1.** Post hoc comparison of Experiment 1.

| Contrast | *t* | *df* | *adj.p* | Cohen's *d* |
| --- | --- | --- | --- | --- |
| RR-OR | 0.22 | 18 | > .999 | 0.05 |
| RR-OO | 3.87 | 18 | .007 | 0.89 |
| RR-RO | 2.91 | 18 | .037 | 0.67 |
| OR-OO | 3.05 | 18 | .035 | 0.70 |
| OR-RO | 2.48 | 18 | .070 | 0.57 |
| OO-RO | 0.51 | 18 | >.999 | 0.12 |

These results revealed that faces whose final facial color was red were more likely to be perceived as angry than faces whose final facial color was consistently neutral. These findings contradict our hypothesis that dynamic changes to facial color affect the recognition of facial expression more than static facial color changes do. Moreover, the last facial color state may be an important cue for the identification of facial expressions rather than the translation of facial color. However, it has been suggested that red attracts more attention and is more strongly bound to objects in memory than green is (Kuhbandner et al., 2015). Therefore, it cannot be conclusively stated from the results of Experiment 1 that a change in facial color has no influence.

We focused on green, which is the color opposite red in the CIE L*a*b* color space. Angry faces with a green facial color (a*−) have been reported to reduce aggression and emotional intensity compared with the original facial color (Thorstenson, McPhetres, et al., 2021; Thorstenson & Pazda, 2021). In other words, by using a green facial complexion, it is possible to enhance the perception of anger in the original facial color, which appears relatively red. This approach allowed us to investigate the effects of differences in facial color on the perception of emotion without relying on red, a color known to strongly influence memory. Therefore, in Experiment 2, we prepared facial stimuli with a decreased red component (a*−) rather than an increased red component. Then, we investigated whether the same results of Experiment 1 would occur when facial stimuli with the original facial color were observed in a relatively redder facial color.

## Experiment 2

The hypothesis suggested that if the identification of facial expression is influenced by dynamic changes to facial color, the PSE for the judgment of facial expressions would differ depending on the presence or absence of changes in color. However, the results of Experiment 1 revealed no significant differences between the change in facial color and no-change conditions. Possible explanations for these results include the influence of the final facial color or the effect of red, which tends to attract attention and may have diminished the effect of dynamic changes to facial color. Therefore, in Experiment 2, we conducted the same task using the color green to determine whether the results of Experiment 1 could be obtained with a reduced influence of the color red. We hypothesized that if dynamic changes to facial color influence the judgment of facial expressions, then reducing the effect of red should lead to a difference between the dynamic color change and no-change conditions. Conversely, if the judgment of facial expressions is influenced by the perception of the final facial color rather than its dynamic change, then results similar to those observed in Experiment 1 are expected.

### Participants

As in Experiment 1, we recruited 22 students (10 women and 12 men; average age 22.4±1.15 years, between 18 and 24 years old) from Toyohashi University of Technology in Experiment 2. All participants had normal color vision, verified by the same color vision test chart used in Experiment 1. They were fully informed about the experiment and consented to participate.

### Stimuli

The facial color states in Experiment 2 were different from those in Experiment 1. The facial color was changed from the original (a*: ±0) to green (−12) or green to the original in the facial color change condition, whereas the color was consistently the original (±0) or consistently green (−12) in the no-change condition (see Figure 4); we defined these conditions as “OG,” “GO,” “OO” and “GG,”, respectively. The other parameters were the same as those used in Experiment 1.

**Figure 4.**
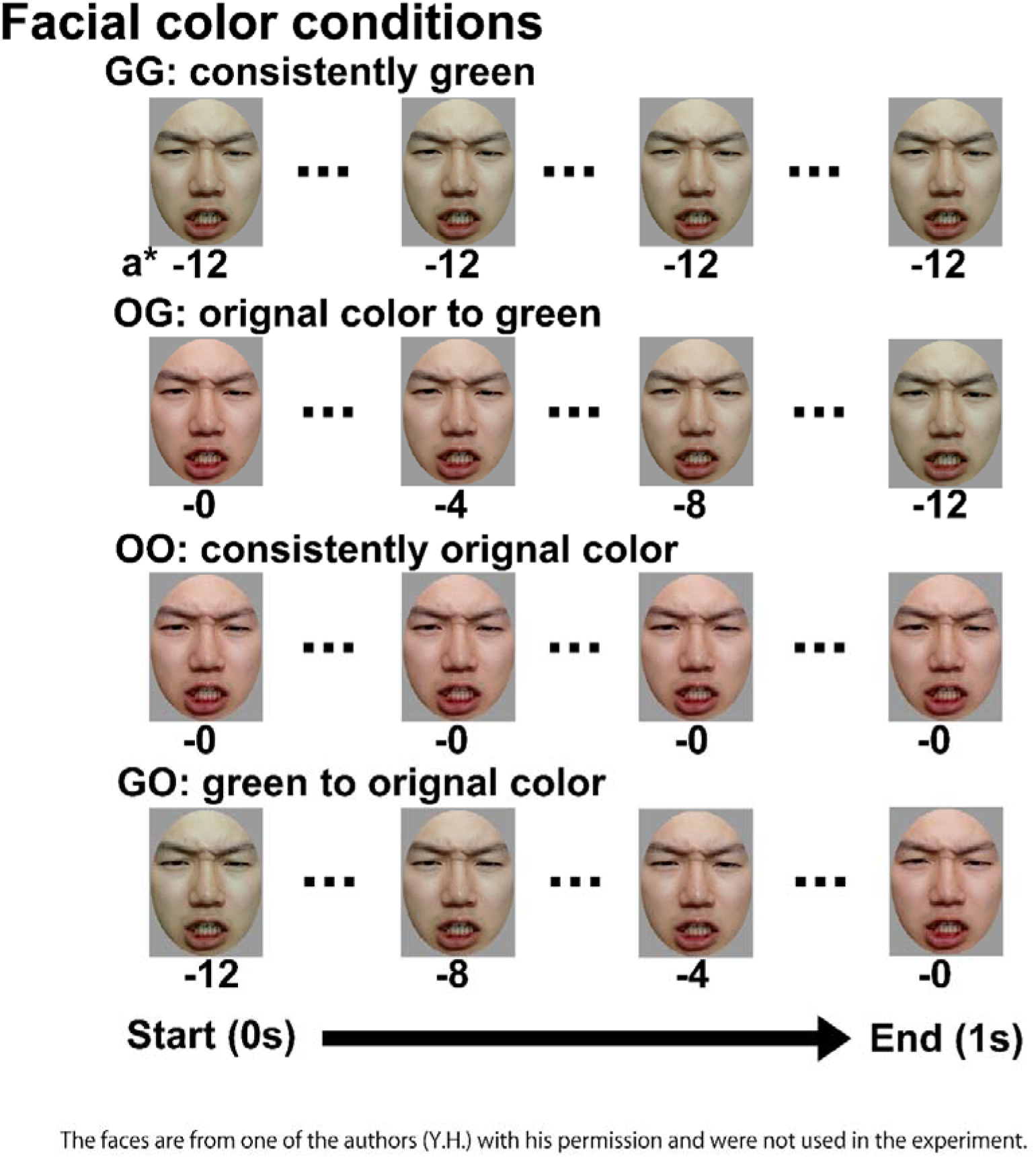
Examples of the image stimuli for each facial color condition in Experiment 2.

### Apparatus

The experimental environment was the same as that of Experiment 1.

### Procedure and task

The procedure was the same as in Experiment 1 except that the presented facial stimuli were different.

### Data analysis

Two participants’ data were excluded from the analysis using the same criteria as in Experiment 1. We confirmed that the participants’ PSEs in each condition were normally distributed. The other parameters were the same as those used in Experiment 1.

### Results and discussion

The average PSEs for each facial color condition are shown in Figure 5. There was a significant main effect of the facial color condition (*F*(1.81,34.39) = 17.23, *p* < .001, *η_p_*^2^ = .476). The statistical results of the post hoc test are shown in Table 2. We observed that the PSEs for OG and GG were significantly greater than those for GO and OO. However, similar to the results of Experiment 1, there were no significant differences between the facial color change and no-change conditions (OG vs. GG and GO vs. OO). Our results suggested that the participants were more likely to perceive the face for which the last facial color was the original color as anger compared to the face for which the last facial color was green.

**Figure 5.**
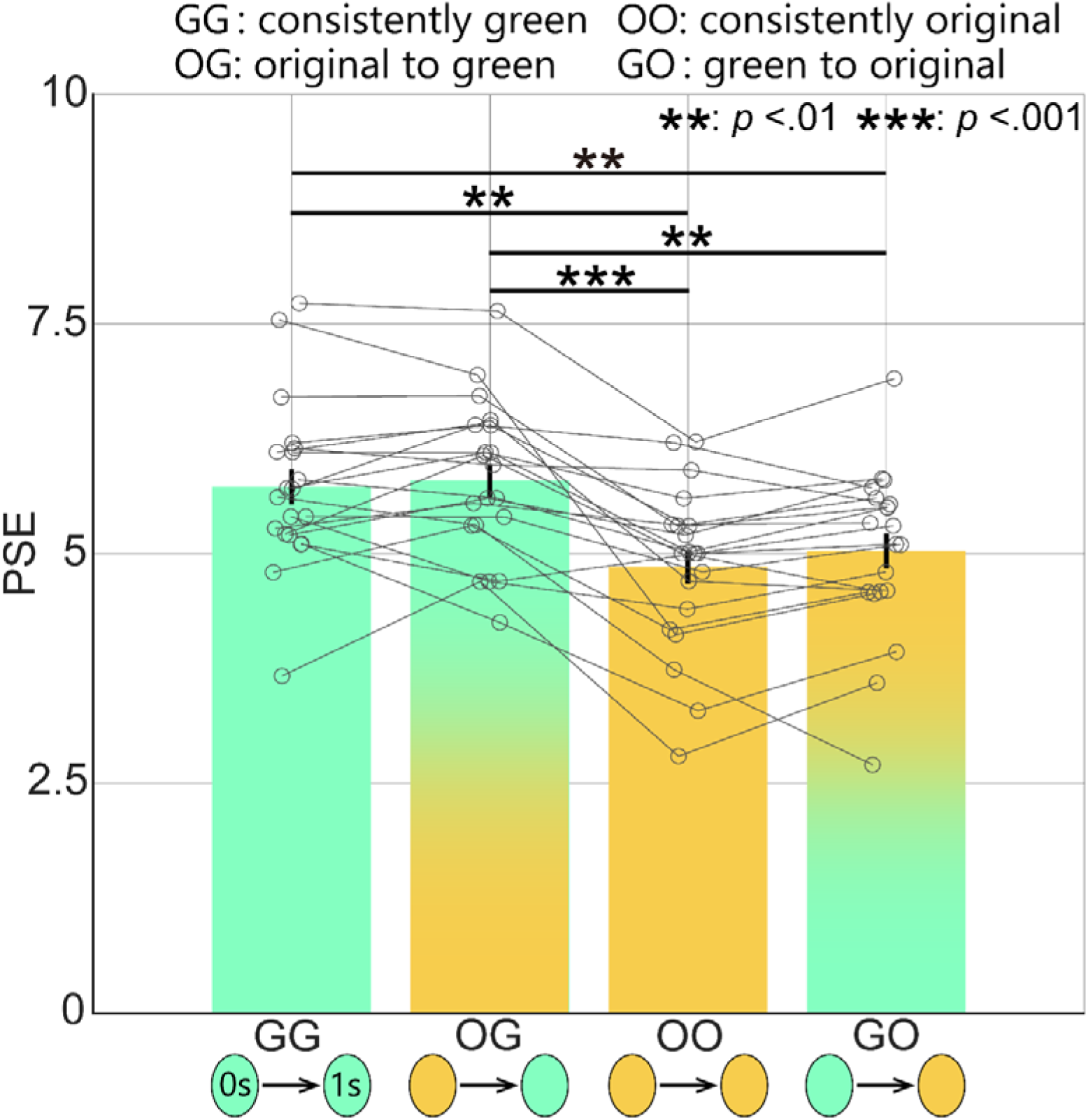
Participants’ mean of PSE in Experiment 2. The other formats were the same as in Figure 3.

**Table 2.** Post hoc comparison of Experiment 2.

| Contrast | $t$ | $df$ | $adj.p$ | Cohen's $d$ |
| --- | --- | --- | --- | --- |
| GG-OG | 0.60 | 19 | .554 | 0.13 |
| GG-OO | 4.56 | 19 | .001 | 1.02 |
| GG-GO | 3.51 | 19 | .007 | 0.79 |
| OG-OO | 5.63 | 19 | <.001 | 1.26 |
| OG-GO | 4.26 | 19 | .002 | 0.95 |
| OO-GO | 1.69 | 19 | .214 | 0.38 |

Our results indicated that the participants were more likely to perceive the face for which the last facial color was the original color as anger, compared to the face for which the last facial color was green. This result suggested that the judgment of facial expression is influenced by the perception of the final facial color rather than its dynamic change.

## General Discussion

In this study, a facial expression judgment task was performed in which different facial color conditions were used to investigate the effects of dynamic changes to facial color on the identification of facial expressions. The results of Experiments 1 and 2 revealed no significant differences between the dynamic change and static facial color conditions. However, a relatively reddish final facial color was likely to be perceived as an angry face. Humans’ retrospective evaluation of experiences and memories is heavily influenced by the peak (most intense) moment and the end (final) moment rather than the overall average or duration(Alaybek et al., 2022; Fredrickson & Kahneman, 1993; Horwitz et al., 2024; Hsu et al., 2018; Kahneman et al., 1993). Therefore, the participants may have been influenced more by the final facial color than by the changes in facial color in the facial expression judgment task. Our results suggest that the identification of facial expressions is affected by the ultimate color rather than the processing of changes to facial color. Previous studies have reported that memory of facial color varies depending on facial expressions. In particular, the memory of the color of an angry face shifts toward a red-yellow color (Hasegawa et al., 2024; Thorstenson, Pazda, et al., 2021). Our finding that facial color at the peak or end of memory influences the recognition of facial expressions may explain these phenomena.

In this study, the perception of anger increased with a reddish facial color (Experiment 1) and decreased with a greenish facial color (Experiment 2). These results are similar to those of previous studies showing that the relationship between facial expression and color modulates human perception and response (Hasegawa et al., 2024, 2025; Kato et al., 2022; Minami et al., 2018; Nakajima et al., 2017; Nguyen et al., 2023; Peromaa & Olkkonen, 2019; Thorstenson, McPhetres, et al., 2021; Thorstenson & Pazda, 2021). These findings are also consistent with those of previous studies in which reddish faces increased the categorization of anger, whereas a green facial color decreased it (Nakajima et al., 2017; Thorstenson, McPhetres, et al., 2021). Our results support the idea that a red facial color increases people’s perception of anger. In addition, it has been reported that reddish angry faces increase the perception of emotional intensity, aggression, and dominance, whereas greenish angry faces decrease these perceptions (Thorstenson, McPhetres, et al., 2021; Thorstenson & Pazda, 2021). The results of this study can be attributed to these changes in the recognition of angry faces due to facial color.

In our experiments, the perception of anger tended to decrease when the facial color shifted dynamically in the opposite color direction to red (RO: red to original condition in Experiment 1; OG: original to green condition in Experiment 2). This result suggests that the judgment of facial expression is influenced by the last facial color even when a red facial stimulus, which is more likely to be perceived as an angry face, is initially presented. While previous studies have dynamically increased the colors associated with emotions (e.g., Thorstenson, Pazda, et al., 2021), our findings on the effects of dynamically decreasing those colors may provide new insights into the relationship between the recognition of facial expressions and color. However, there was a relatively large change in facial color because the change in this study was a*+12 or −12 within 1 second, whereas in the experiment by Thorstenson et al. (2021), it was a* and b* +5 or −5 within 1 second (Thorstenson, Pazda, et al., 2021). Therefore, it is reasonable to consider the results of this study to be limited in scope.

In this study, we found that the perception of anger was lower for greenish faces than for the original facial color (green vs. original; Experiment 2). Most previous studies have compared differences in the perception of facial expressions using red, the original facial color, and blue (red vs. original vs. blue; Nakajima et al., 2017) or have compared red and green (red vs. green) or red, the original facial color, and green (red vs. original vs. green; (Thorstenson & Pazda, 2021). A few studies have explored the use of relative redness or compared the original color and green. Our findings suggest that the judgement of facial expressions is influenced by even relatively reddish facial colors, such as combinations of original and green colors. These findings provide new evidence that supports perceptual responses in the relationship between facial expression and color.

The limitations of this study that should be addressed in future research are as follows. First, because this study focused on the differences between dynamic changes to facial color and static facial color, the presence or absence of perceived changes to facial color was prioritized as a key factor. On the basis of previous studies, we adopted simple parameters for the duration and amount of changes in facial color during the experiment with a linear change of a*+12 in one second (Nakajima et al., 2017; Thorstenson, Pazda, et al., 2021). However, facial color changes in nature are neither uniform nor linear (Jimenez et al., 2010; Kikuchi et al., 2015; MORETTI et al., 1959; Zonios et al., 2001). It has been reported that when individuals express anger, facial blood flow increases, causing the face to appear redder (Drummond, 1999; Drummond et al., 2001; Kreibig, 2010). However, this color change is not influenced by a single hue alone (not only a* channel) but rather by the colors inherent in substances such as hemoglobin and melanin, which are included in blood flow (Edwards & Duntley, 1939; Kikuchi et al., 2015; Zonios et al., 2001). In addition, due to variations in the thickness and blood vessel distribution of facial skin, facial color changes do not necessarily occur with the spatial uniformity observed in the experiments of this study (MORETTI et al., 1959; Tsumura et al., 1999, 2003). Experiments that use facial color manipulation related to components of hemoglobin or melanin (e.g., Kato et al., 2022) as well as experiments that utilize nonlinear facial color changes should be conducted in the future.

Second, the effects of color may have been strong. As previously explained, the parameters of a* in this experiment were set on the basis of prior research by Nakajima et al. (2017), but the amount of color change was relatively large. Thus, it is possible that the effect of dynamic facial color changes was overshadowed by the saliency of the color.

Third, we did not research the individual characteristics of the participants. It has been reported that the ability to recognize faces and facial expressions as well as attention biases toward specific emotions vary depending on individual characteristics such as trait anxiety (Surcinelli et al., 2006; Telzer et al., 2008). Therefore, the results of this study might reflect the influence of individual characteristics. Further examination of variations in the effects arising from differences in these characteristics is warranted.

Finally, all participants in both experiments were Japanese or familiar with Japanese culture. In addition, the facial stimuli were Japanese faces. Hence, the findings of this study should be interpreted as being based on phenomena observed under specific conditions, and their validity is confined to particular populations because it has been suggested that the judgment of facial color is based on the average skin tone observed in daily life (Shimakura & Sakata, 2022).

## Conclusions

We conducted two experiments using facial expression stimuli with different facial color presentation conditions (Experiment 1: red and original color; Experiment 2: green and original color) to investigate whether dynamic changes in facial color influence the recognition of facial expressions. The results revealed that there were no significant differences between dynamic facial color changes and consistently identical facial color conditions. These findings suggest that the identification of facial expressions is affected by the final perception of facial coloration rather than by the process of facial color change. Our findings offer new insights into the relationship between facial expression recognition and facial color.

## Data availability

All data except for the stimuli are publicly available at the https://osf.io/p7ey9/?view_only=81aa2032dbfd4c4ba2bd7741c834f9d7.

## Author contributions

M.S., Y.H. and T.M. designed the research; M.S., Y.H. and T.M. provided experimental code and conducted the analysis; M.S. Y.H., H.T., S.N. and T.M. wrote the paper; Y.H., H.T., S.N. and T.M. were responsible for funding acquisition.

## Acknowledgments

This work was supported by Grants-in-Aid for Scientific Research (KAKENHI) from the Japan Society for the Promotion of Science (Grant Numbers JP22K1789 to H.T., JP20H05956 to S.N., JP20H04273 to T.M., and JP23KK0183 to T.M.), the Supporting Pioneering Researchers in Transformative Research (SPRING) program from the Japan Science and Technology Agency (Grant Number JPMJSP2171 to Y.H.), and the student fellowship program for the Leading Graduate School at Toyohashi University of Technology to Y.H.

